# Accumulation of ^137^Cs in Insects - Herpetobiont Integuments

**DOI:** 10.1101/853242

**Authors:** D. Monoshyn, L. Rudchenko, T. Shupova, V. Gaychenko

## Abstract

The study of the radioactivity of insects - herpetobiont elytra that belong to different trophic groups shown the presence of incorporated ^137^Cs, there were concentrated about 20% of total animal radioactivity. The comparison of the radionuclide comparison was done in both dried and fresh samples. Radioactive contamination of integument comes mainly from the terminal phase of metamorphosis (at the ninth phase).

## Introduction

In nearest future, the questions of intaking, accumulation, biological transformation, and redistribution of the radioactive compounds by domestic and wild animals are not going to escape careful look of radiobiology and radioecology. Especially important place in this question is taken by the study of the radioactive contamination of invertebrates, especially of insects. These animals are characterized by a relatively short life cycle, high fertility, and biomass. They have taken almost all biological niches, which makes them ones that have important (or even main) influence in the transformation of matter and energy at the natural or half-natural ecosystems. It also allows us to use them as the object of ecological and, especially, radioecological monitoring [1].

## Materials and methods

The insect with the different trophic specializations, collected from Drevlyany radioecological reserve (Zytomirska oblast, Ukraine, the density of radioactive contamination of soil is more than 25 kBq/m^2^) at 2017 served as material for the study. The activity of recently sampled and dry material was analyzed. To determine the activity of ^137^Cs in integuments elytra were studied.

Collected samples were shredded, measurements were done using Gamma - spectrometry equipped with Ge(Li) half conducting detector GEM-30185, GMX (EG&G ORTEC) and multichannel analyzer (ADCAM-300, USA, IN-1200, France). To perform the analysis, the measuring vessels “Denta” (shape of a frustum, 3.3cm high, 6.3 and 7.3 cm diameter of bases). Measurement has been done in the Institute of Agricultural Radiology NULES Ukraine.

The insects - herpetobionts of different ecological groups were studied: phytophagous (Zabrus tenebrioides Goeze, 1777, Melolontha melolontha L. 1758), predators (Calosoma inquisitor L. 1758, Calosoma auropunctata Herbst, 1874), detritophages (Nicrophorus vespillo L. 1758), and coprophagous (Geotrupes stercorarius L. 175).

## Results and discussion

Results of the measurements showed that in all cases integument is characterized by a relatively high level of radioactive contamination (Table 1). A high observational error may be explained by the low weight of samples.

**Table 1.** Activity of insect integuments.

| Specie | Trophic specialization | Mass of the sample, g | Activity, Bq/kg | Observation error, % |
| --- | --- | --- | --- | --- |
| *Calosoma inquisitor* | Predator | 5,914 | 540 | 15 |
| *Calosoma auropunctata* | Predator | 24,21 | 3335 | 10 |
| *Zabrus tenebrioides* | Phytophagous | 4,992 | 600 | 22 |
| *Nicrophorus vespillo* | Necrophagous | 4,813 | 630 | 15 |
| *Geotrupes stercorarius* | Coprophagous | 10,587 | 1510 | 12 |

As previous researches showed, redistribution of 1^137^Cs in the trophic chain (from phytophagous and up to coprophagous) for insects is following the same pattern that known for other animals - increasing of concentration along with the moving to the higher point on the trophic chain [4, 5, 7].

Since dry entomological material was used for the research, fresh samples of Melolontha melolontha L. in situ (whole body and elytra only) w were used as a control and to compare the contamination level (Table 2). As can be seen from the table, the activity of “fresh” Melolontha melolontha and their elytra shows that insect integument contains about 26% of total animal radioactivity. The suggestion that other studied insect species following the same or at least similar relation was taken as a working hypothesis.

**Table 2.**
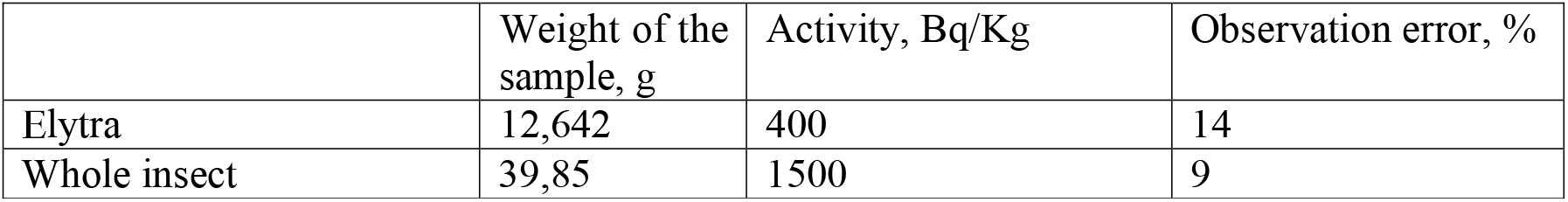
Activity of whole *Melolontha melolontha* and their elytra.

On the modern state of radioactive contamination of the forests of Ukrainian Woodland, the main place where radionuclides are accumulating is forest floor and topsoil [6], which are ecological niches for the majority of the species of insects - herpetobiont. The main way of radioincludes income is the trophic one since larvas and imagos intake contaminated fodder and accumulate biologically significant radionuclides, such as ^137^Cs. Radioactivity of insects integuments requires more attention since it is not yet described in the literature.

Radioactive contamination of insect integument occurs while it is forming at the larva phase, which is characterized by high activity of potassium [8, 9], whose chemical analysis is ^137^Cs. A relatively low amount of radionuclide comes to an animal at the imago stage, but, according to authors, it concentrates on fat and tissues.

Taking to account significant biomass of entomofauna and its important role in the migration of radionuclides at the natural ecosystems, peculiar properties of redistribution of ^137^Cs by the insects - herpetobionts, as well as its partition in the organs and tissues of animals require future investigation.

## List of sources

1. Gaychenko V.A. Faunistic complex as an object of radioecological monitoring // Nuclear physics and energy. 14, 2 (2013) 295–298.

2. Romanenko V.N. Fundamentals of the comparative physiology of invertebrates. (Tomsk: Tomsk State University 2013) 224 p.

3. Comparative physiology of animals. Ed. L. Prosser. T. 1. (M .: 1977) 608 p.

4. V. Gaychenko, D. Monoshyn. Redistribution of ^137^Cs in trophic chains of insectsherpetobions. // Science Newsletter NUBIP of Ukraine. Seriya “Biology, Biotechnology, Ecology”. 287 (2018) 7–14.

5. V.A. Gaychenko, V.D. Naumova. Migratsiya ^137^Cs of the Chornobilsky similarity according to the trophic lancet of the pastoral type. // Mat. scientific-practical conference up to 120 riches NUBIP of Ukraine. (2018) p. 70 – 73.

6. Krasnov V.P., Shelest Z.M., Kurbet T. V, Boiko O. L. ^137^Cs redistribution in time in wet bory and sugrudy solls in Forests of Ukrainian Polissia // Nuclear physics and atomic energy. –17 (2016) 394–399.

7. Simonova L.I., Gaychenko V.A. Biogenic ^137^Cs migration in trophic lanceugs // Ukrainian Radiological Journal 2 (2009) 218–220

8. Arthur M. Jungreis, William R. Harvey Role of active potassium transport by integumentary epithelium in secretion of larval-pupal molding fluid during silkmoth development // J. exp. Biol. (1975), 6a, P. 357–366 (https://www.sciencedirect.com/science/article/abs/pii/S0065280608600521)

9. Arthur M. Jungreis Physiology of Moulting in Insects // Advances in Insect Physiology. Volume 14, 1979, P. 109–183 (https://www.semanticscholar.org/paper/Role-of-active-potassium-transport-by-integumentary-Jungreis-Harvey/c80d20ed043755a9dbf7fbddb9dd87b4eddcecea)

